# Genetic landscape of populations along the Silk Road: admixture and migration patterns

**DOI:** 10.1101/011759

**Authors:** Massimo Mezzavilla, Diego Vozzi, Nicola Pirastu, Giorgia Girotto, Pio d’Adamo, Paolo Gasparini, Vincenza Colonna

**Author notes:** Corresponding Author Vincenza Colonna Institute of Genetics and Biophysics ‘A. Buzzati-Traverso’ National Research Council (CNR) via Pietro Castellino 111 - 80131 Napoli – Italy.

## Abstract

**Background:** The ancient Silk Road has been a trading route between Europe and Central Asia from the 2nd century BCE to the 15th century CE. While most populations on this route have been characterized, the genetic background of others remains poorly understood, and little is known about past migration patterns. The scientific expedition “Marco Polo” has recently collected genetic and phenotypic data in six regions (Georgia, Armenia, Azerbaijan, Uzbekistan, Kazakhstan, Tajikistan) along the Silk Road to study the genetics of a number of phenotypes.

**Results:** We characterized the genetic structure of these populations within a worldwide context. We observed a West-East subdivision albeit the existence of a genetic component shared within Central Asia and nearby populations from Europe and Near East. We observed a contribution of up to 50% from Europe and Asia to most of the populations that have been analyzed. The contribution from Asia dates back to ~25 generations and is limited to the Eastern Silk Road. Time and direction of this contribution are consistent with the Mongolian expansion era.

**Conclusions:** We clarified the genetic structure of six populations from Central Asia and suggested a complex pattern of gene flow among them. We provided a map of migration events in time and space and we quantified exchanges among populations. Altogether these novel findings will support the future studies aimed at understanding the genetics of the phenotypes that have been collected during the Marco Polo campaign, they will provide insights into the history of these populations, and they will be useful to reconstruct the developments and events that have shaped modern Eurasians genomes.

## BACKGROUND

The ancient Silk Road has been a trading route for several centuries in the past (2^nd^ century BCE - 15^th^ century CE) serving as a main connection between Europe and Asia. The route traverses a geographical region that was central during the human expansion from Africa [1, 2], however its role during the stages of human evolution and its recent genetic history have not been fully clarified. At any rate, the complicated demographic history of the Silk Road populations must have left a trace on the patterns of genetic variation in a very unique way. Previous studies based on uniparental markers revealed extensive admixture between Europeans and eastern Asians in central Asia, increased genetic diversity and correlation with linguistic patterns [3–5]. Similar results were observed in a study based on genomic data from twenty-seven microsatellites genotyped in 767 individuals from Uzbekistan Kyrgyzstan and Tajikistan [6]. This study reported a pattern of diffused admixture and migration among populations in the area, resulting in great genetic diversity correlated with patterns of linguistic diversity, as it is often the case [7]. Recently a more comprehensive study disentangled the different levels of gene flow in worldwide populations but did not include a number of populations form Central Asia, for which migration patterns are still not fully understood [8].

With the aim of studying the genetics of several phenotypes (e.g. food preferences, hearing and taste [9–12]), the Marco Polo scientific campaign, collected in 2010 genetic and phenotypic information of 411 individuals from six countries dwelling along the Silk Road (Georgia, Armenia, Azerbaijan, Uzbekistan, Kazakhstan and Tajikistan). Some of them have not been extensively described in population genetics literature, and the work we present here aims to investigate for the first time the population structures and the admixture patterns by using hundreds of thousands of genome-wide markers and 53 other worldwide populations as a reference for comparison. We confirmed results of previously mentioned studies, and extended the analysis to quantify admixture proportions and migration contributions. Here, we are providing insightful information about the different levels of gene flow in these populations, highlighting the differences in population history and genetic structure.

## RESULTS AND DISCUSSION

### Great genetic diversity and east-west blocks in the Silk Road populations

We compared genetic data at 299,899 single nucleotide variants for 441 individuals from six populations along the Silk Road (abbreviated in SR from now on) with 943 individuals from 53 populations worldwide from the Human Genome Diversity Project (HGDP) panel[13]. To make the discussion of the results easier we grouped HGDP populations by their geographical area, and we will refer to the populations form Central Asia newly genotyped in this study as Silk Road, even if they are not exhaustively representative of the Silk Road (Table S1).

First, we explored global patterns of genetic variation in our study populations. In the principal component analysis (PCA) of all 53 populations (Figure S1), both first and second components distribute SR populations along the Asia-Europe gradient, and close to the nearby populations from Central Asia. SR populations, especially Uzbekistan and Kazakhstan, span over a region of the plot only slightly smaller in size than the one separating Europe from East Asia, suggesting great genetic diversity in these populations. ADMIXTURE analysis[14] on the same set of populations (Figure S2), revealed, under the most likely hypothesis of ten clusters, very little or no contribution from Africa, Oceania and America and thus, we repeated analyses after removing these populations. The resulting PCA plots (Figures 1A and Figure S3) show two different patterns for the Western and the Eastern SR. The Western SR (WSR) including Armenia, Georgia, and Azerbaijan shows, proximity with Europe and the Near East, while the Eastern Silk Road (ESR) including Uzbekistan and Tajikistan and Kazakhstan shows a proximity generally closer to Asia, except for a number of individuals from Uzbekistan and Kazakhstan who are closer to Europe and the WSR. This pattern does not reflect the sampling strategy (rural communities *versus* cities, see Subjects and Methods) but rather true genetic diversity; indeed samples closer to Europeans are from the same sampling locations of samples in the other block. Admixture analysis on the same reduced set of populations, revealed more details. The most likely hypothesis of seven clusters (Figure 1B, see results for different number of clusters in Figure S4) shows that, overall, populations of the SR have a similar amount of European ancestry (red in Figure 1B) suggesting a common genetic legacy between Europe and the SR. An exception is found for some individuals in Uzbekistan and Kazakhstan where this component is >50%; these individuals cluster with Europe in the PCA. There are instead two opposite gradients in the WSR and the ESR for components shared with Near East and Central-South/East Asia. In the WSR we observe predominance of the Near Eastern component (pink and light green in Figure 1B), whereas in the ESR there is a predominance of Asian components (shades of blue in Figure 1B). Notably the dark blue component, which is diffused within the ESR, is predominant in the Kalash isolate [15] (Figure S4), suggesting a common ancestral origin. The overall picture suggests that, beside a shared genetic pattern, there are two blocks of populations, and the ESR populations tend to be more admixed than the WSR ones.

**Figure 1.**
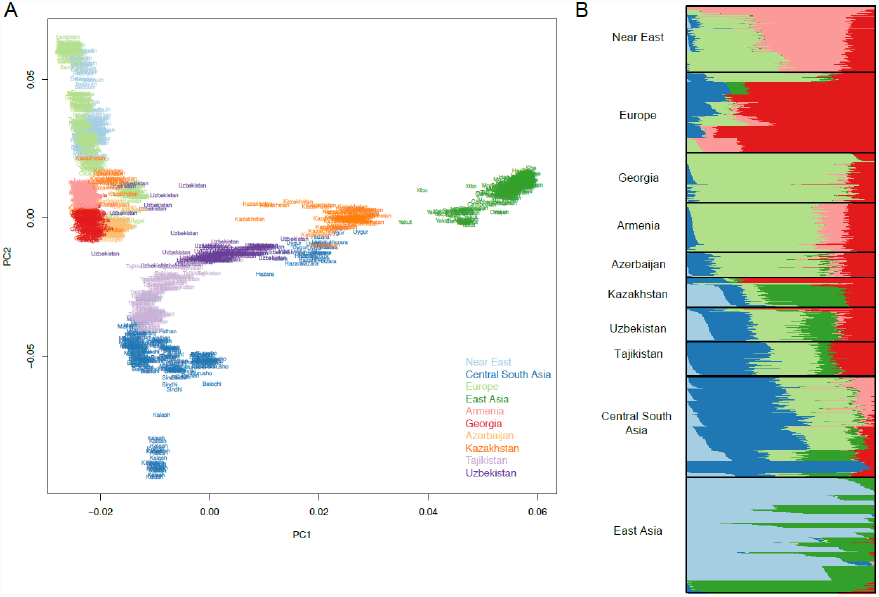
Population genetic structure. A. Principal component analysis of populations from the Silk Road newly genotyped in this study (SR) with Near Eastern, European, Central-South and East Asian populations. The color scheme reflect geographical regions, the first and second axes explain the 26 % and 8.7 % of the variance, respectively. SR’s populations form two distinct blocks. ***B.*** Model based admixture analysis in the hypothesis of seven clusters. SR’s populations have in general similar amount of European ancestry and there are two opposite gradients in Western and Eastern SR for components shared with Near East and Central-South/East Asia. Both in ***A*** and ***B*** a number of individuals from Uzbekistan and Kazakhstan are closer to Europe and WSR.

The hypothesis of the two blocks, is further corroborated by the positions that the SR populations occupy in the tree of the populations’ divergence times estimates, as described in [16] (Figure 2) with the WSR close to Europeans, and the ESR between Central-South Asia and East Asia, with a large split time difference.

**Figure 2.**
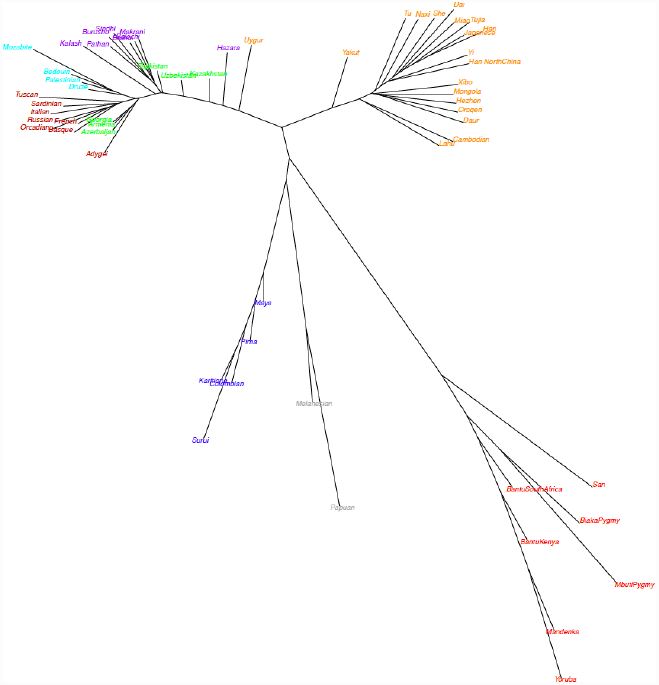
Populations divergence times. Unrooted neighbor joining tree of 50^th^ percentile of time of divergence (measured in years) between populations.

Next we explored the history of isolation and consanguinity in our set of populations. When calculating inbreeding coefficient, we did not observe particularly high values for the SR populations (Figure S5). We evaluated the prevalence of stretches of homozygous regions (ROH) by comparing the cumulative distribution of the total length of ROH per individual in each population or group of populations, assuming a minimum ROH length of 2 Mb. Among the Silk Road populations, Tajiks have the longest stretches of homozygosity (Figure 3), higher than Europeans and East Asians (Figure S6) and only lower than other founder populations, e.g. Surui in America, Papuan in Oceania and Kalash in Central-South Asia. In 80% of the Tajiks, the stretches of homozygosity sum up to 100 Mb whereas for other populations this value is below 30%. Because the length and the extension of ROH reflect the effective population size (Ne)[17, 18], we confirmed our findings by evaluating N_e_ as described in [19–21] Tajikistan has one of the lowest N_e_ among SR populations, with a declining trend in the last 10,000 years (Figure S7). However, for all SR populations, long-term estimates N_e_ is comparable to other reference populations (Figure S8), suggesting high level of genetic variation in that area.

**Figure 3.**
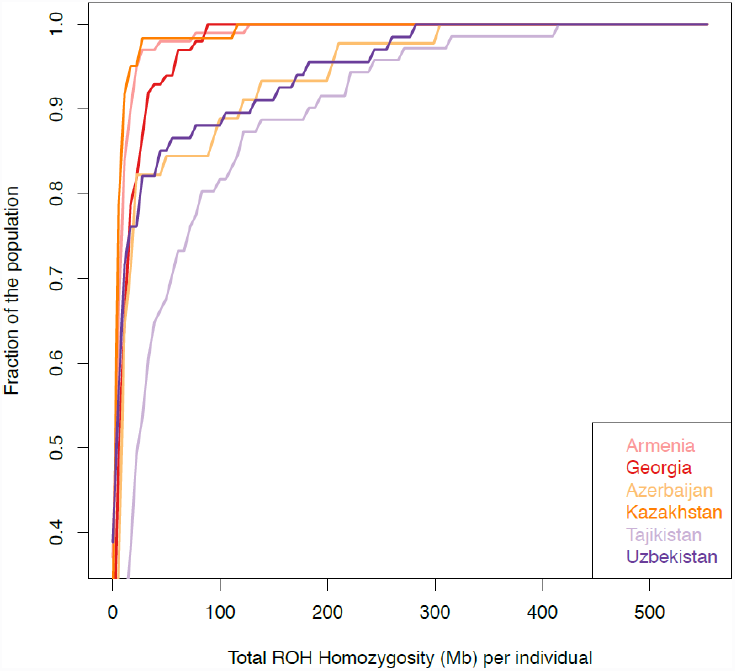
Cumulative distribution of runs of homozygosity. The x-axis indicates the total homozygosity in the genome in Megabases (Mb). Minimum ROH length is set to 2 Mb.

### Migrations patterns revealed unidirectional gene flow from East Asia to Eastern Silk Road

We used two approaches to evaluate the role and effects of gene flow among populations in this study. Firstly, we assessed admixture for each population from any pair of other two populations using the three-population test for admixture [22, 23]. All possible combinations of populations were performed, but we reported only results with Z-score <-5 in Table S2. Overall, Z-score results indicate little or no gene flow in WSR, except for a number of contributions to Azerbaijan (mainly for Armenia and East Asia, see Table S2), including contributions from American populations that we interpret more as a shared origin rather than an introgression. On the contrary, many migration events in the ESR are observed. Uzbekistan and Kazakhstan show the highest level of mixing in accordance with results from ADMIXTURE and PCA analyses (Figure 1), whereas Tajikistan received contributions mainly from Central-South Asian populations. We concluded that, as for admixture, we observed clearly different patterns of migrations in the WSR compared to the ESR, with more migration events taking place in the ESR, consistently to the degree of admixture that we inferred.

In order to look in more detail at migrations, we built a tree of all populations through ancestry graph analysis [23] (Figure 4). The tree structure recapitulates the known (existing) relationships among these populations, and highlights the great complexity of this geographical area, as previously shown by the neighbor joining tree, based on the time of divergence (Figure 2). The tree built with all the populations from HGDP reaches the maximum variance explained (99.7%) with the lowest standard error of residuals, with nine migration edges (Figure 4A). The tree shows the WSR populations compacted near Europe and the Near East, whereas the ESR populations are scattered (Figure 4A). Tajikistan falls with Central-South Asia; Kazakhstan with East Asia with a migration from Adygei; Uzbekistan is the target of a migration from the branch which connects East Asian and Americans.

**Figure 4.**
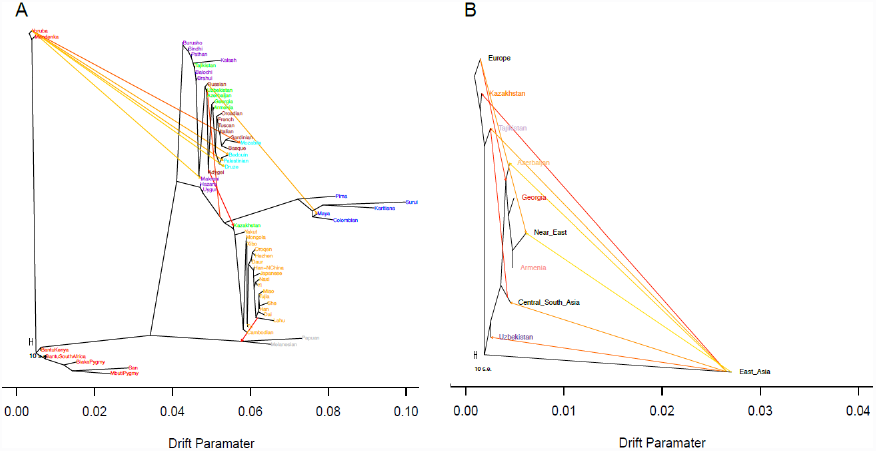
Migration patterns. A. Tree with nine migration edges connecting all the population in this study. ***B.*** Tree without America, Africa and Oceania (nine migration edges) suggesting unidirectional gene flow from East Asia to Eastern Silk Road populations and Azerbaijan.

To focus only on migrations relevant to the Silk Road populations, we built a tree removing Africa, Oceania and America (Figure 4B). The resulting tree with 9 migration edges explains 99% of the variance that was observed. Six migration edges connect East Asia with the SR. Indeed, with this analysis we pinpoint the gene flow from East Asia to the ESR and we estimate that the fraction of alleles that East Asia donated as source to other populations, ranges from 0.14 to 0.44, major recipient being Kazakhstan (Table 1). Tajikistan also received a consistent contribution (0.42) form Central-South Asia, thus highlighting the different history of this population within the ESR. As expected, we found gene flow from Europe to the WSR (0.39 towards Georgia and Azerbaijan) and a very low contribution of East Asia to Azerbaijan (0.05), but not to the other WSR populations. With these analyses we identified and quantified gene flow within the SR.

**Table 1.** Migration edges estimates from Treemix

| Source | Target | Allele sharing<br>(standard deviation)<br>in % |
| --- | --- | --- |
| East Asia | Kazakhstan | 44 (0.8) |
| East Asia | Uzbekistan | 27 (0.1) |
| East Asia | Tajikistan | 14 (0.2) |
| Central-South Asia | Tajikistan | 42 (2.3) |
| Europe | Georgia / Azerbaijan /<br>Near East / Armenia | 39 (1) |
| East Asia | Azerbaijan | 5.1 (0.2) |
| Europe | Near East | 18 (0.9) |
| East Asia | Central-South Asia | 18 (0.05) |
| East Asia | Near East | 1.5 (0.3) |

Our results suggest unidirectional gene flow from East Asia to the ESR, and apparently this gene flow never reached the WSR, except for Azerbaijan, that was affected in a minor mode.

### To what extent populations admixed?

Having assessed the general pattern of genetic relationships between populations, we next quantified admixture between populations. We first estimated the fraction of reciprocal contributions using ALDER [24] with 1 reference curve. In Figure 5 we summarized fractions of contributions from source populations to target. We estimated that the WSR received contributions from Europe and Near East in a measure of 40-50%. Armenia shows the highest level of Near East ancestry (48%) and Azerbaijan is the only population receiving significant contributions from the ESR (Kazakhstan 18%, Uzbekistan 35%). Among the ESR populations and in general, Uzbekistan and Kazakhstan are the more admixed populations with contributions of respectively 47% and 48% from Europe, and 49% and 47% from Central-South Asia. For these two populations, admixture within the SR is also high, ranging from 40% to 80%, and they also have the highest score of admixture (~80%) between themselves. Finally, East Asian components are higher in Kazakhstan compared to Uzbekistan and Tajikistan, as reported previously [25]. These results are consistent with ADMIXTURE analyses that we illustrated in a previous section. We also corroborated these observations by evaluating haplotype sharing among populations through chromosome painting. We used SupportMix [26] to assign haplotypes of individuals in SR populations to reference populations in the HGDP panel. In Figure S9 we summarized the fraction of loci in target SR populations assigned to reference HGDP populations and we observe that these are consistent with admixture trends estimated from ALDER and ADMIXTURE. Details of SupportMix analyses are reported in Table S3.

**Figure 5.**
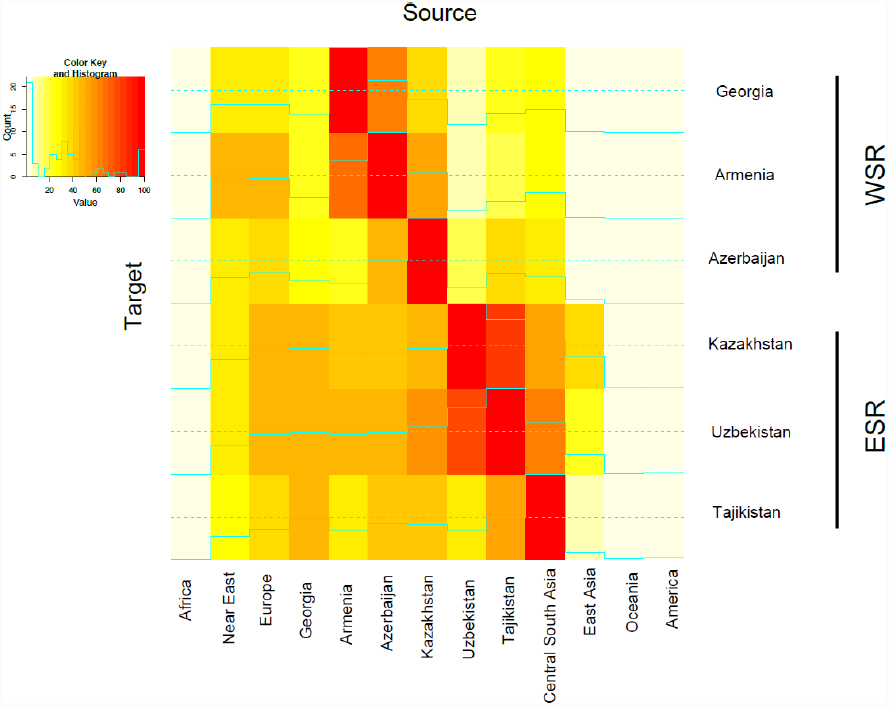
Admixture between pairs of populations. Heat map showing admixture contributions of source populations to targets inferred from linkage disequilibrium patterns. Rows are the target populations, columns are the source populations. Admixture with the same population is set at 100% and coded in red. Full lines indicate exact values and dashed line mark the 50%.

To sum up, our data demonstrated poor admixture between the WSR and the ESR: only Azerbaijan seems to have a connection with the ESR. Within the WSR there is a non-negligible contribution of the Near East to Azerbaijan, but not to Georgia and Armenia, despite the three populations clustering together in the PCA analysis. Within the ESR, Tajikistan is generally less admixed than the other two populations. This could reflect a history of isolation and related low gene flow within the ESR, as suggested by the pattern of long stretches of homozygosity (Figure 3). Finally, Uzbekistan and Kazakhstan are the more admixed, both between themselves, and with respect to others, as hypothesized from PCA analyses (Figure 1).

### When did admixture events take place?

We inferred the time in generations during which main migration events took place from patterns of linkage disequilibrium decay [24], and reported results that remain significant after Bonferroni’s correction in Table S4. Within the WSR, we observed quite ancient events: a flow from Georgia and the Near East to Armenia ~210-220 generations ago (solid arrows in Figure 6), and a flow from the ESR and East Asia to Georgia ~60 generations ago (dot and line arrows in Figure 6). Among recent events (~25 generations ago, dashed lines in Figure 6) in the WSR, we observed contributions of East Asia to Azerbaijan (consistent with the results in the previous section); ESR populations instead are characterized by several introgressions from Central-South and East Asia.

**Figure 6.**
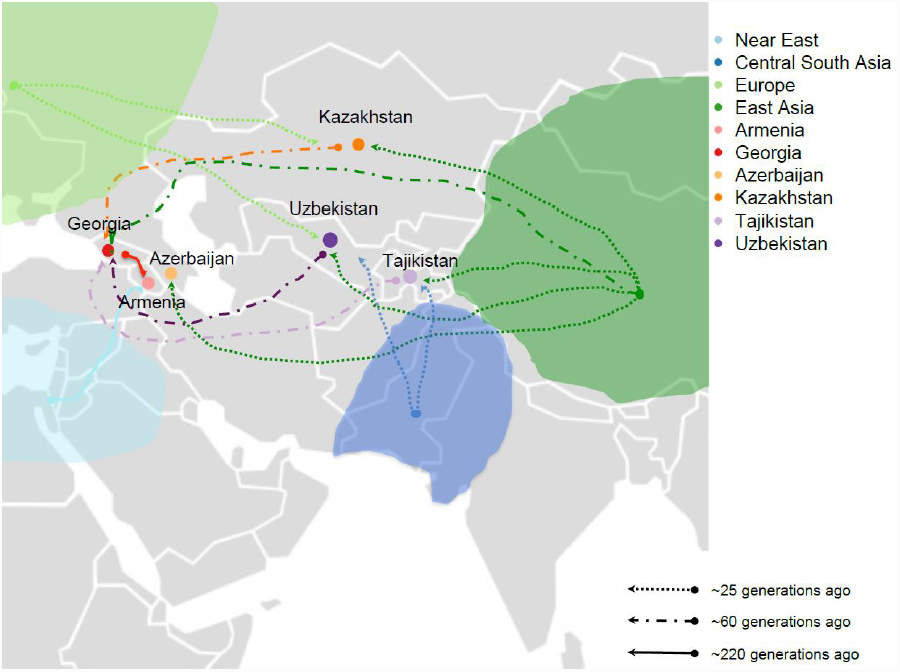
Estimates of dates of migration events. The map is showing the geographic location of the six Caucasian and Central Asian populations sampled during the “Marco Polo” scientific expedition and newly genotyped in this study. Arrows on the map provide information on migration events detected in this study. Arrows paths, origin and destination should be interpreted with caution in that they reflect a population trend rather than a specific geographic location.

The overall picture is of a flux around 25 generations ago from Central-South and East Asia towards the West that only reached Azerbaijan through the WSR, and this is consistent with the gradient of contributions of Central-South and East Asia described in Figure 5. Considering a generation time of 25 years, this scenario is compatible with the Mongolian Empire expansion, as can be observed in [8]. Finally, Tajikistan shows an additiona event, even more ancient, involving Central-South Asia ~40 generations ago, which could explain the high level of haplotype sharing obtained using SupportMix compared to Uzbekistan and Kazakhstan, thus confirming a previous study based on Y chromosome [27].

Having demonstrated that indeed admixture took place among the populations that were taken into account for this study, and having assessed directions of migrations, thanks to these analyses, we now add extra information on times during which these events might have occurred. Time estimates should be interpreted with caution, but in combination with historical information might be useful to understand past events.

## CONCLUSIONS

We provided a detailed description of some demographic events that shaped genetic diversity in some of the populations along the Silk Road. We observed high genetic heterogeneity in patterns of admixture and genetic structure. We identified a main subdivision in Western and Eastern Silk Road and an East-West gradient of East Asia contribution reaching only Azerbaijan within the West and taking place ~25 generations ago, a time compatible with the Mongolian expansion. A fine-scale resolution of these admixture events will be possible with whole genome sequencing of these populations. Our findings provide a catalogue of genetic variation that will help to better understand the phenotypic variations that we previously described in these countries [9–12] [10, 28] and provide information on time and extent of past migration events that can be integrated with historical studies.

## SUBJECTS AND METHODS

### Ethic Statement

All communities belong to the Terra Madre organization (www.terramadre.org). Approval for information collection and processing has been obtained by the Ethical committee of the Maternal and Child Health Institute IRCCS-Burlo Garofolo Hospital (Trieste, Italy). All individuals signed an appropriate consent form (written in their local language) after they were instructed about the project, and all samples were completely anonymous.

In addition to the Italian Ethical committee approval we obtained the approval from the local Ethical committee and from the National Council on Bioethics in Georgia. For other countries where Ethical Committee were not present at the time of sampling we received authorization through official direct approval letters by National Authorities (Ministry of Research or Ministry of Health).

### Samples and genotypes

Subjects in this study have been enrolled as part of the scientific campaign “Marco Polo”, aimed at collecting social, genotype and phenotype information from populations of central Asia and Caucasus along the Silk Road. Populations in this study originate from West to East: Georgia, Armenia, Azerbaijan, Uzbekistan, Kazakhstan, Tajikistan and for the sake of simplicity we will collectively refer to them as the Silk Road populations, even if they are not fully representative of the entire range of countries along the Silk Road (Table S1). Samples of saliva were collected in these countries from healthy donors from rural communities (Georgia, Azerbajan, Armenia, Tajikistan or form both rural communities and cities (Uzbekistan, Kahzakstan). DNA was extracted from samples that were collected, and it was used to determine genotypes at 624,851 single nucleotide polymorphic sites (SNPs) as described in a previous study [28]. For all subsequent analyses we used Plink v.1.07 [29] unless differently specified. Genomic coordinates refer to the GRCh37/hg19 version of the human genome sequence. Relatedness among pairs of individuals within populations was calculated as the proportion of genome that is identical by descent (pi_hat in Plink) and one random individual per pair related above first or second cousins (pi_hat > 0.25) was kept for subsequent analyses. After removing individuals that failed genotype quality controls and/or were related, the final sample consisted of 441 individuals. As reference populations, we used publicly available data from HGDP [13], after removing elated subjects as suggested in[30]. We merged genotype information from the Silk Road populations with HGDP genotypes at the 22 autosomal chromosomes filtering for minor allele frequency >1%, and genotyping success rate >97%. After quality controls, the merged data set consisted of 299,899 SNPs genotypes.

### Population structure, effective population size and time of divergence estimates

Shared ancestry between populations was evaluated using ADMIXTURE v 1.22 and the number of cluster that better represent the data was established by cross-validation as described in [14]. Each ADMIXTURE run was replicated 5 time using different random seeds. Principal Component Analysis (PCA) analysis was performed using EIGENSOFT [31], considering the difference in sample size between our populations and HGDP ones. Our samples from the Silk Road were projected using the axis obtained from the HGDP populations. PCA and ADMIXTURE were performed on a linkage disequilibrium (LD) pruned dataset (r^2^<0.4), consisting of 186267 SNPs.

Runs of homozygosity (ROH) were calculated using 299,899 SNPs. A ROH is defined as a chunk of genome ≥ 2000 kb long, containing at least 100 SNPs, with SNP density >= 1 per 50 kb. Within a ROH one heterozygous SNP and five missing calls are allowed. Two consecutive ROHs are considered as a single unit if their distance is <= 1 Mb. Coefficient of inbreeding was estimated using the function *–het* implemented in PLINK. Demography and long-term effective population size were estimated by using linkage disequilibrium information as described in [16]. Since this method requires the genetic map position to be known, we used HapMap recombination map to assign a genetic position. Time of divergence between populations was estimated as described in [16] implemented in the R package NeON [32], a unrooted neighbour joining tree was built using the R package ape [33]. The generation time considered is of 25 years per generation.

### Time of admixture and gene flow

Treemix v1.12 [23] was used to estimate maximum likelihood tree of our populations. Two different dataset were used: the first one with all the population and the second with population merged as continental groups without Africa, America and Oceania (see Table S1 for group labeling). Zero to 12 migration edges were modeled, using blocks of 500 SNPs and the *-se* option was used to calculate standard errors. The tree was rooted with Yoruba when using the dataset with all the populations. Migration edges were added until the model explained 99% of the variance and also the migration edges were significant. Migration rate was estimated from Treemix run for the best tree selected. **A** Three-population test [22] was used to assess evidence of admixture in each of the six Silk Road population, values with a Z-score of less than -5 were collected. Level and time of admixture events were estimated using ALDER v.1.0.3 [24], considering the high number of possible comparisons and to avoid bias in allele frequencies due to small samples, we merged each reference population in continental groups.

Significance of the admixture was assessed after Bonferroni’s correction and consisted in the decay of linkage disequilibrium. In addition the level of admixture was assessed for each population using as reference for the population of the Silk Road only one population at the time. As an additional approach, we applied a machine learning method, SupportMix [26], to estimate genome-wide ancestry. This method allows to infer loci-specific genomic ancestry when analyzing different reference populations. SupportMix was run on phased datasets using a sliding window of 500 Kb for each chromosome and each individual.The mean percentage of the assigned chunk of genomes to each continental group was collected as well. Phasing was obtained using BEAGLE [34]. A mean value of assigned loci was calculated for each population of the Silk Road.

## ADDITIONAL FILES

1) mezzavilla_supp_tab.xls

## LIST OF ABBREVIATION

SR: Silk Road, N_e_: effective population size, ROH: Runs of homozygosity. WSR: Western Silk Road, ESR: Eastern Silk Road

## AUTHORS’ CONTRIBUTION

MM, VC, PG conceived and designed the study. GG, PG, NP, DV, PDA collected samples, MM analyzed the data. MM, VC wrote the paper. All authors read and approved the final manuscript.

## ACKNOWLEDGMENTS

We wish to thank all the subjects from the populations of Georgia, Armenia, Azerbaijan, Kazakhstan, Uzbekistan and Tajikistan who donated saliva samples for DNA analyses. We also wish to thank Irene Dall’Ara and Guido Barbujani for their contributions in the early stage of the project and Qasim Ayub and Chris Tyler-Smith for comments and suggestions on the manuscript. Finally, we thank two anonymous reviewers for helpful comments on the manuscript. The study was funded by the sponsors of the scientific expedition Marcopolo 2010, and by Ministry of Health RC 35/09 to PG The funders had no role in study design, data collection and analysis, decision to publish, or preparation of the manuscript.

---

**Figure S1.**
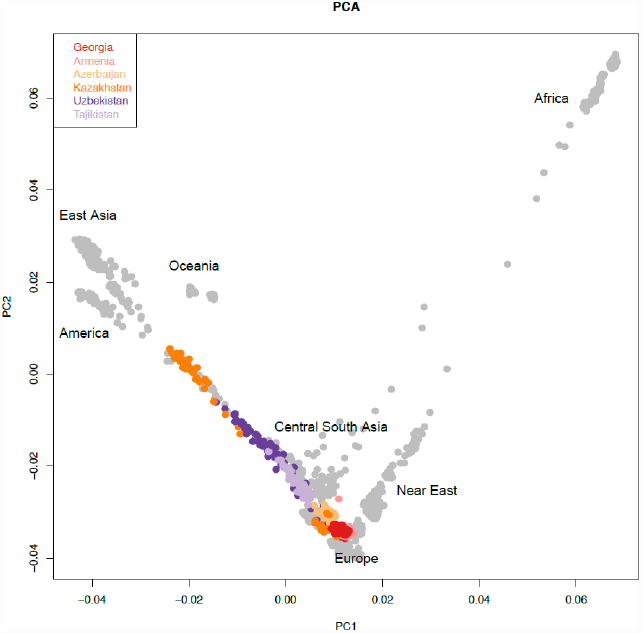
Principal component analysis of populations from the Silk Road newly genotyped in this study (in color) with populations in the Human Genome Diversity Project panel (in grey). Silk Road populations, and especially Uzbekistan and Kazakhstan, span over a large region of the plot, suggesting great genetic diversity among individuals. The first axis explain the 5.6% of the variance, the second axis 4.3 % of the variance

**Figure S2.**
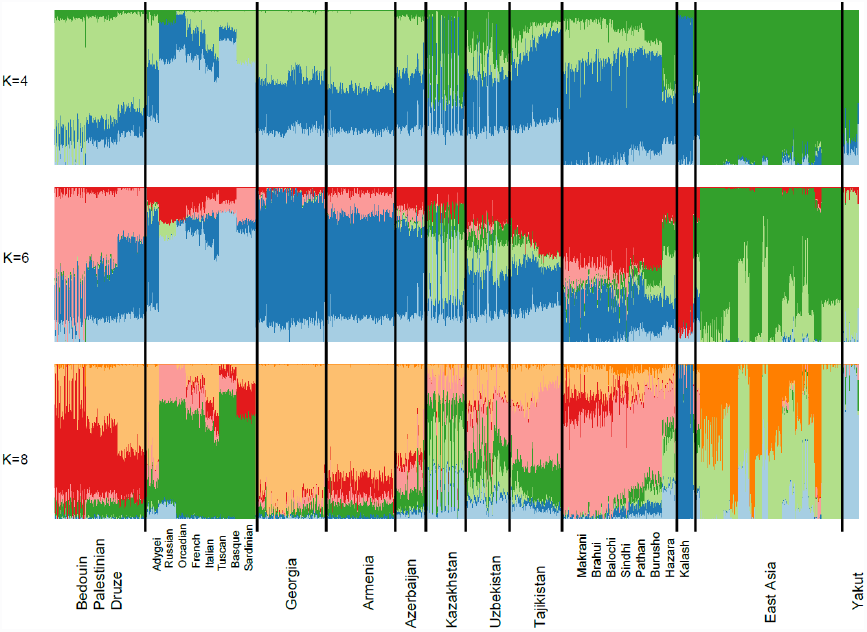
Admixture plots for the hypotheses of 8, 9 and 10 clusters (K). The most likely hypothesis (K=10) shows little or no contribution from Africa, America and Oceania to populations newly genotyped in this study.

**Figure S3.**
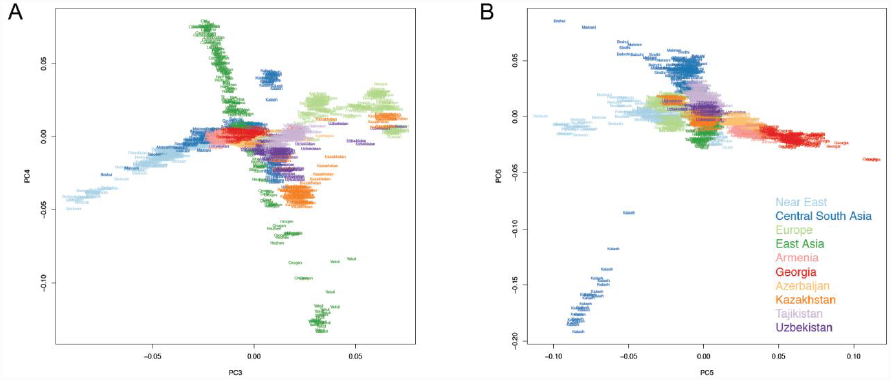
Principal Component Analysis of a subset of worldwide populations. Representation of components 3 to 6

**Figure S5.**
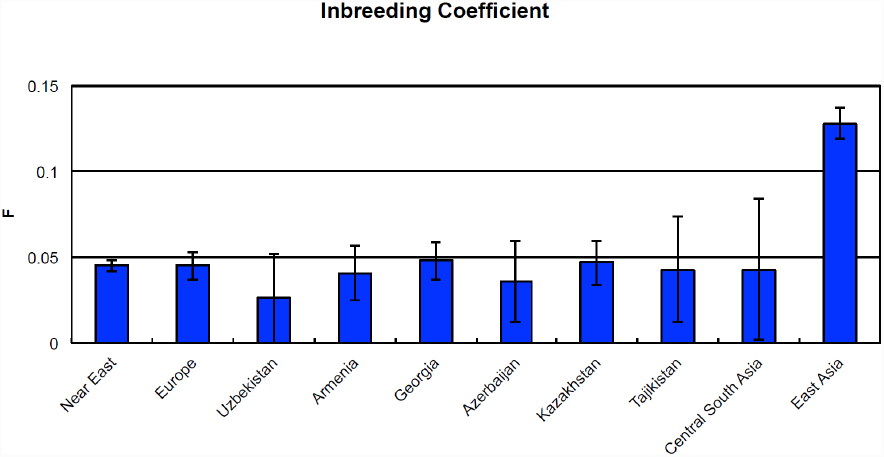
Inbreeding coefficients. Averages of individuals inbreeding coefficients (y-bars represent standard deviation) per population or group of populations.

**Figure S6.**
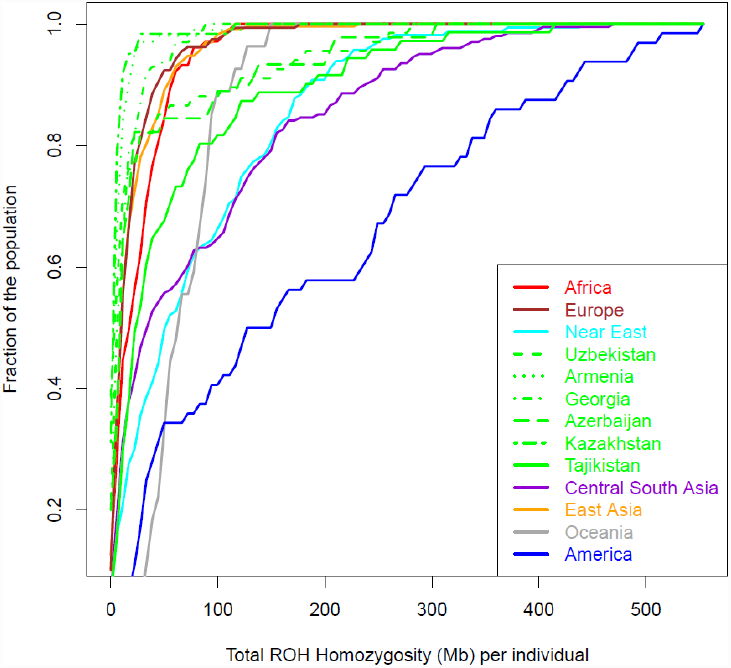
Cumulative distribution of runs of homozygosity. The x-axis indicate the total homozygosity in the genome in Megabases (Mb). We set the minimum ROH length to 2 Mb.

**Figure S7.**
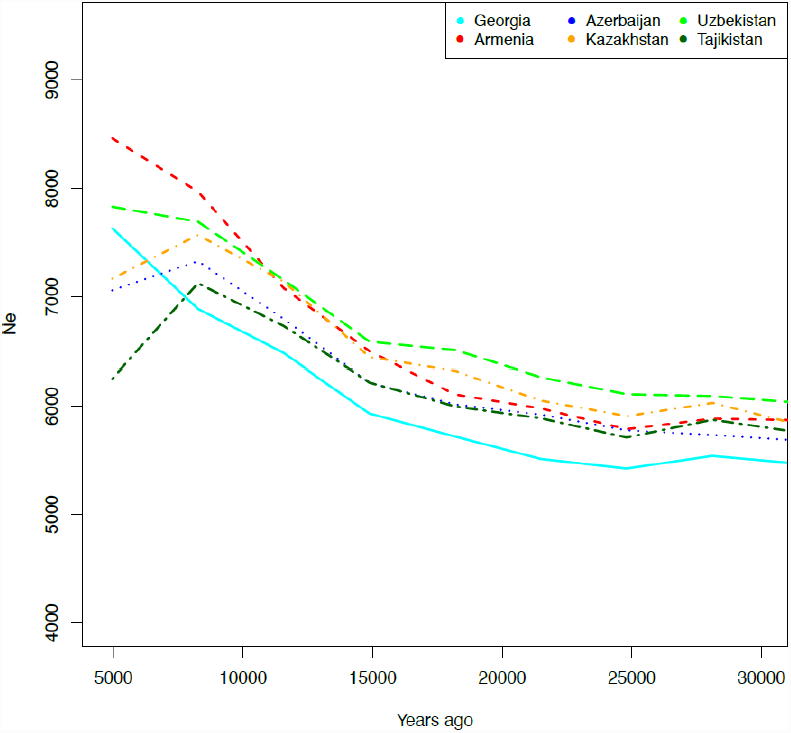
Effective population size through time. The y-axis represent the effective population size (Ne), the x-axis the time in the past. Generation time is 25 years

**Figure S8.**
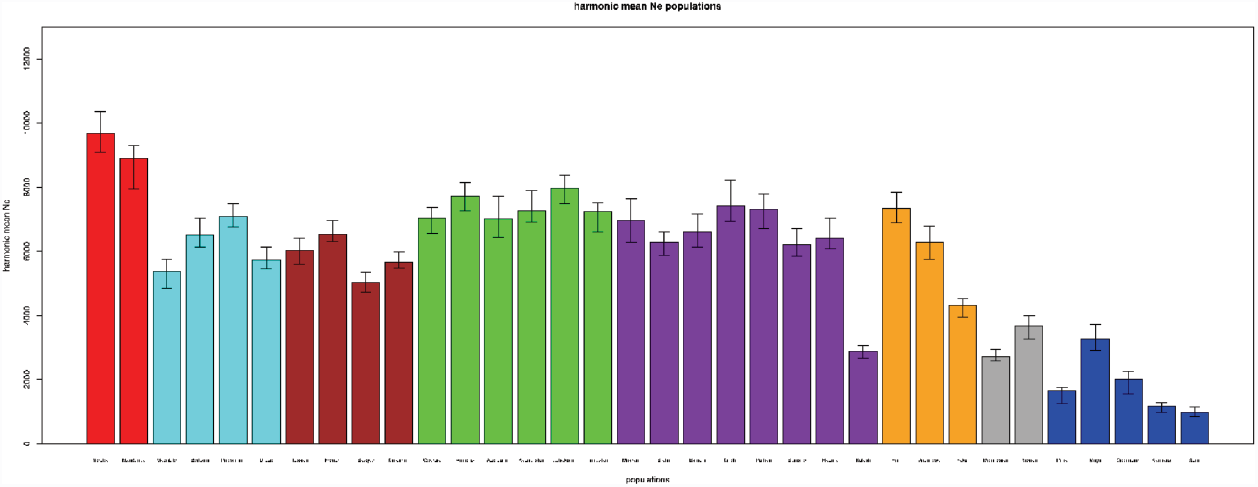
Long term estimate of the effective population size.

**Figure S9.**
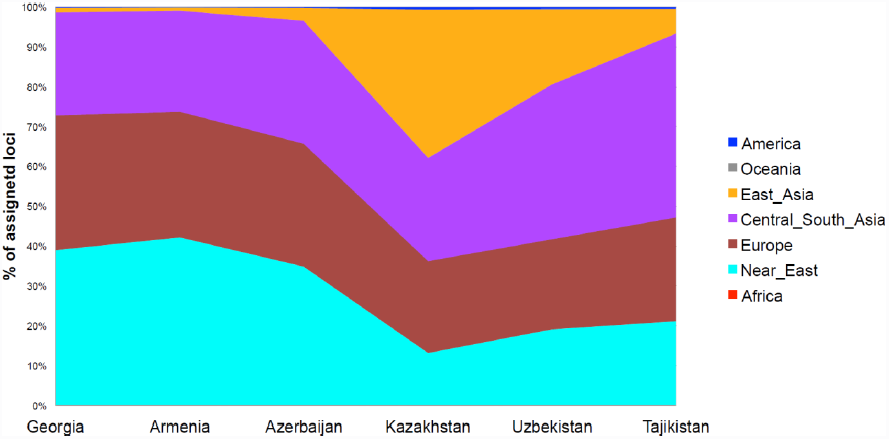
Proportions of admixture between pairs of populations inferred from haplotype sharing. On the y-axis the mean percentage (estimated across 22 autosomes) of genomic segment assigned to different continental groups (see colors in the legend) per each population on the x-axis.

***Table S1. Population description and sample size.***

***Table S2 Three population test statistic.*** Only results with Z-score < -5 are reported and only when populations belong to different continental groups.

***Table S3. SupportMix mean percentage of assigned loci across al 22chromosomes.*** Rows are the references, columns are the targets.

***Table S4. Time of admixture estimate using ALDER.***

